# CD209L/L-SIGN and CD209/DC-SIGN act as receptors for SARS-CoV-2

**DOI:** 10.1101/2020.06.22.165803

**Authors:** Razie Amraei, Wenqing Yin, Marc A. Napoleon, Ellen L. Suder, Jacob Berrigan, Qing Zhao, Judith Olejnik, Kevin Brown Chandler, Chaoshuang Xia, Jared Feldman, Blake M. Hauser, Timothy M. Caradonna, Aaron G. Schmidt, Suryaram Gummuluru, Elke Muhlberger, Vipul Chitalia, Catherine E. Costello, Nader Rahimi

## Abstract

As the COVID-19 pandemic continues to spread, investigating the processes underlying the interactions between SARS-CoV-2 and its hosts is of high importance. Here, we report the identification of CD209L/L-SIGN and the related protein CD209/DC-SIGN as receptors capable of mediating SARS-CoV-2 entry into human cells. Immunofluorescence staining of human tissues revealed prominent expression of CD209L in the lung and kidney epithelium and endothelium. Multiple biochemical assays using a purified recombinant SARS-CoV-2 spike receptor binding domain (S-RBD) or S1 encompassing both NTB and RBD and ectopically expressed CD209L and CD209 revealed that CD209L and CD209 interact with S-RBD. CD209L contains two *N-*glycosylation sequons, at sites N92 and N361, but we determined that only site N92 is occupied. Removal of the *N*-glycosylation at this site enhances the binding of S-RBD with CD209L. CD209L also interacts with ACE2, suggesting a role for heterodimerization of CD209L and ACE2 in SARS-CoV-2 entry and infection in cell types where both are present. Furthermore, we demonstrate that human endothelial cells are permissive to SARS-CoV-2 infection and interference with CD209L activity by knockdown strategy or with soluble CD209L inhibits virus entry. Our observations demonstrate that CD209L and CD209 serve as alternative receptors for SARS-CoV-2 in disease-relevant cell types, including the vascular system. This property is particularly important in tissues where ACE2 has low expression or is absent, and may have implications for antiviral drug development.

## Introduction

The outbreak of coronavirus disease 2019 (COVID-19), which is caused by severe acute respiratory syndrome-coronavirus-2 (SARS-CoV-2), constitutes a serious ongoing threat to global public health and has generated a major worldwide socio-economic impact ^1–3^. Morbidity and mortality of SARS-CoV-2 are associated with acute respiratory distress syndrome (ARDS) and other complications such as coagulopathy, thrombosis and multi-organ failure in COVID-19 patients ^3–6^. Although the role of the vascular system, particularly endothelial cells, in the pathogenesis of COVID-19 remains largely unknown, emerging evidence suggests that SARS-CoV-2 directly attacks the vascular system ^7–9^. Severe endothelial injury, vascular thrombosis with micro-angiopathy, occlusion of alveolar capillaries, and angiogenesis were distinctively observed in lung autopsies of COVID-19 patients ^10^, underscoring the critical importance of the vasculature system in the pathogenesis of COVID-19.

Human angiotensin-converting enzyme 2 (ACE2) is known to interact with the surface spike (S) protein of SARS and also acts as an entry receptor for SARS-CoV-2 ^11–13^. While it was previously reported that ACE2 is widely expressed in the lung, vascular system and other organs^14^, recent studies demonstrated that ACE2 is expressed at very low levels and only in a small subset of lung epithelial cells ^15^ and low-to-undetectable levels in endothelial cells^16^, suggesting that SARS-CoV-2 entry into and infection of certain human cells may be occurring via alternative receptors or a combination of multiple receptors and/or enhancers. Consistent with this idea, Neuropilin receptors ^17–18^, CD147/Basigin ^19^ and heparin sulfate ^20^ are reported to facilitate SARS-CoV-2 entry. Neuropilin receptors are highly expressed in endothelial and neuronal cells, and play major roles in vascular endothelial growth factor (VEGF)-dependent angiogenesis and semaphorin-dependent axon guidance ^21^. CD147 is expressed in erythrocytes ^22–23^ and endothelial cells of the brain and acts as a receptor for plasmodium ^22, 24^. In addition to ACE2, alternative receptors that function as points of entry have been reported for other coronaviruses, such as human NL-63 and SARS-CoV ^25^. These include CD209L (also known as L-SIGN) and CD209 (also known as DC-SIGN) ^26–29^.

CD209L and CD209 are members of the C-type lectin superfamily and are implicated as mediators of viral pathogenesis ^25, 30^. While CD209L is highly expressed in human type II alveolar cells and the endothelial cells of lung, liver and lymph nodes ^31–32^, CD209 is primarily expressed in dendritic cells and tissue-resident macrophages, including alveolar macrophages ^33^, dermal macrophages ^34^, and peripheral blood mononuclear cells ^35–36^. Despite their differential expression profiles, CD209L and CD209 share 79.5% amino acid sequence homology. The most distinguishing region of CD209L and CD209 is the C-type lectin domain (CRD), which functions as a calcium-dependent glycan-recognition domain ^37^. It is thought that the highly conserved EPN motif (Glu-Pro-Asn) on the CRD domain is responsible for the recognition of mannose, fucose- or galactose-containing structures^38^. However, despite a high degree of homology of the amino acid residues in the CRD of CD209L and CD209, there is evidence for differential recognition of oligosaccharide structures by these receptors. For example, CD209L appears to prefer high mannose oligosaccharides but not complex glycans, especially those containing antennary fucose epitopes, as LewisX (LeX), whereas CD209 binds to fucose and LeX ^39–40^.

In this manuscript, we demonstrate that the receptor-binding domain (RBD) of the SARS-CoV-2 S protein binds to CD209L and CD209, mediating SARS-CoV-2 entry. CD209L is expressed in human endothelial cells and mediates endothelial cell adhesion, capillary tube formation and sprouting. CD209L contains two sequons (NXT/S, X≠ P) that provide the potential for *N*-linked glycosylation at sites N92 and N361. We determined that only the N92 site is occupied and that high mannose glycans are present at this site. Removal of *N*-glycans from the cell surface enhanced the binding of S-RBD with CD209L. We further show that CD209L interacts with ACE2, in tissues where both are present, and thereby propose a role for heterodimerization of CD209L and ACE2 in virus entry and infection. These findings suggest that CD209L and CD209 represent novel potential therapeutic targets against COVID-19 and have implications for antiviral drug design.

## Results

### CD209L is expressed in human lung epithelial and endothelial cells and regulates the angiogenic properties of endothelial cells

To investigate the potential involvement of CD209L and CD209 in COVID-19, we first examined expression of CD209L in human tissues from SARS-CoV-2 target organs, which include lung endothelial and epithelial cells, renal vessels, tubules, and glomeruli, and temporal artery, via immunofluorescence staining. Lung tissue was co-stained with antibodies against CD209L and MUC1, the latter serving as a marker for type II alveolar cells ^41–42^, and was examined using laser microscopy. Isotope-labeled antibody served as a control and showed negligible staining of the lung tissue (**S. Figure 1A**). The specificity of CD209L antibody was validated in cell culture in HUVEC-TERT cells. Knockdown of CD209L eliminated the immunoreactivity of CD209L antibody in Western blotting (**S. Figure 1B**). Our results showed prominent expression of CD209L in the MUC1 positive alveolar cells (**Figure 1A**). We also observed expression of CD209L in pulmonary capillaries (**Figure 1B**), endothelium of the small and medium sized temporal artery (**Figure 1C**) and renal arterioles (**Figure 1D**). CD31 served as a marker of endothelial cells (**Figure 1B, C, D**). Moreover, we show that CD209L is expressed in renal proximal tubular epithelial cells, which were marked by aquaporin1 (**S. Figure 2A**). However, CD209L was not observed in the glomerular capillaries and minimal expression was noted in the mesangial area, probably in the infiltrating immune cells (**S. Figure 2B**). The results demonstrate the prominent expression of CD209L in type II alveolar cells and pulmonary endothelium, as well as renal vessels and renal tubular cells which are also potential target cells of SARS-CoV-2.

**Figure 1.**
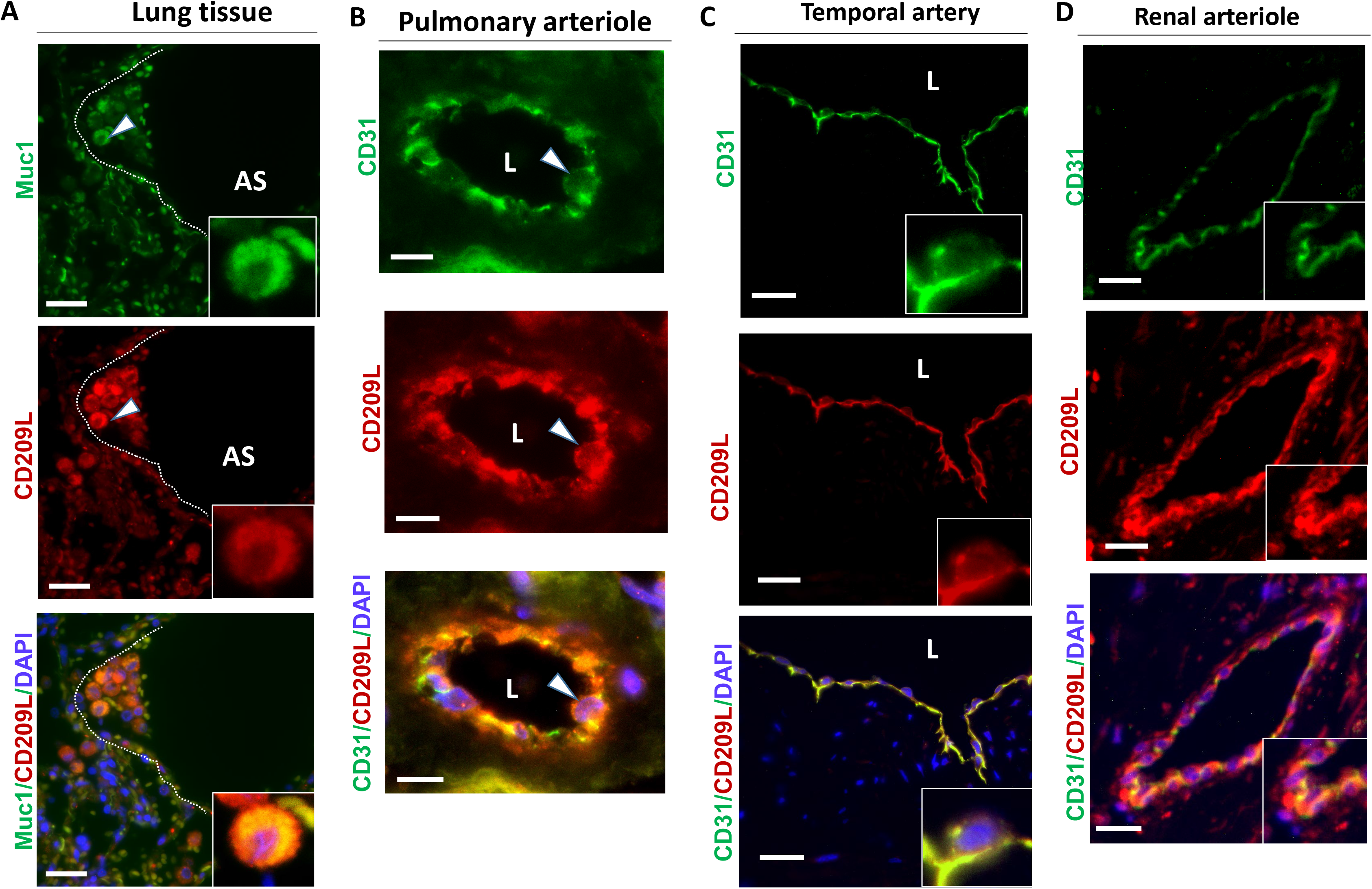
CD209L is expressed in lung, and renal epithelial and endothelial cells: PFA fixed human lung, renal and temporal arteriole tissues were subjected to immunofluorescence staining. Lung tissue stained with anti-MUC1, anti-CD31, and anti-CD209L antibodies. (**A**) Type II alveoli epithelial cells of alveoli were positive for CD209L (red) and MUC1 (green). (**B**) Pulmonary arteriole endothelial cells were positive for CD31 (green) and CD209L (red). (**C**) Endothelial cells of temporal arteriole were stained with CD209L (green) and CD31 (red). (**D**) Renal endothelial cells were positive for CD209L (red) and CD31 (green). White arrowhead pointing to the alveolar cell and AS= alveolar space and white dotted line corresponds to alveolar septa.

CD209 was expressed only in a subset of type MUC1 positive II alveolar cells (**S. Figure 2A**). Furthermore, unlike CD209L which is highly expressed in endothelial cells (**Figure1C, D**), we did not observe expression of CD209 in the pulmonary or renal arterioles (data not shown). A distinct expression of CD209 was observed in the limited proximal tubular epithelial cells in kidneys (**S. Figure 3B**), but glomerular capillaries were also mostly negative for CD209 (**S. Figure 3C**). Overall, these results suggest that CD209 has a limited expression profile compared to CD209L in the organs examined.

Considering that CD209L is highly expressed in endothelial cells, we examined the potential role of CD209L in regulation of the angiogenic responses of endothelial cells. First, we confirmed expression of CD209L in human umbilical endothelial cells immortalized with telomerase (HUVEC-TERT cells) ^43^ by Western blot analysis (**S. Figure 4A**).However, expression of ACE2 in these cells was very low, and multiple weak protein bands potentially corresponding to ACE2 were only detected after long exposure times (**S. Figure 4B**). Additionally, we compared ACE2 expression in HUVEC-TERT cells to lung carcinoma cell lines, A549 and H1299 cells. ACE2 detected in the cell lysates of A549 and HT1299, but not in the HUVEC-TERT cells (**S. Figure 4B**). To address the functional importance of CD209L in endothelial cells, we knocked down CD209L via shRNA strategy (**S. Figure 4C**) and tested key angiogenic characteristics of endothelial cells, including cell adhesion, capillary tube formation/*in vitro* angiogenesis and cell migration. To determine the role of CD209L in cell adhesion, we generated a Myc-tagged soluble CD209L encompassing the ectodomain of CD209L (sCD209L) (**Figure 2**). We coated 24-well plates with sCD209L and incubated HUVEC-TERT cells expressing control shRNA or CD209L-shRNA. After 30 minutes incubation, un-adhered cells were washed off the plates, cells were fixed and the number adherent cells was quantified. The result showed that depletion of CD209L in HUVEC-TERT cells significantly decreased cell adhesion (**S. Figure 6A**), suggesting that CD209L mediates endothelial cell-cell contact. Next, we subjected these cells to an *in vitro* angiogenesis assay. HUVEC-TERT cells expressing CD209L-shRNA displayed considerably reduced capillary tube formation compared to HUVEC-TERT cells expressing control shRNA (**S. Figure 5B**). Capillary tube formation is a complex dynamic process that involves cell-cell adhesion, cell proliferation and cell migration. Therefore, we examined the effect of knockdown of CD209L in the migration of HUVEC-TERT cells. Knockdown of CD209L resulted in a robust increase in cell migration (**S. Figure 5C**). Given that remodeling of the actin cytoskeleton into filopodia, formation lamellipodia (*i.e.*, cytoplasmic protrusions that contain a thick cortical network of actin filaments), and stress fiber assembly plays vital roles in endothelial cell migration ^44^, we investigated actin stress fiber formation by staining of the cells with phalloidin. Consistent with the observed effect of knockdown of CD209L in cell migration, silencing of CD209L also increased cytoplasmic protrusions at the leading edge of HUVEC-TERT cells (**S. Figure 6**). These data demonstrate that CD209L serves important roles in the regulation of angiogenic properties of endothelial cells.

**Figure 2:**
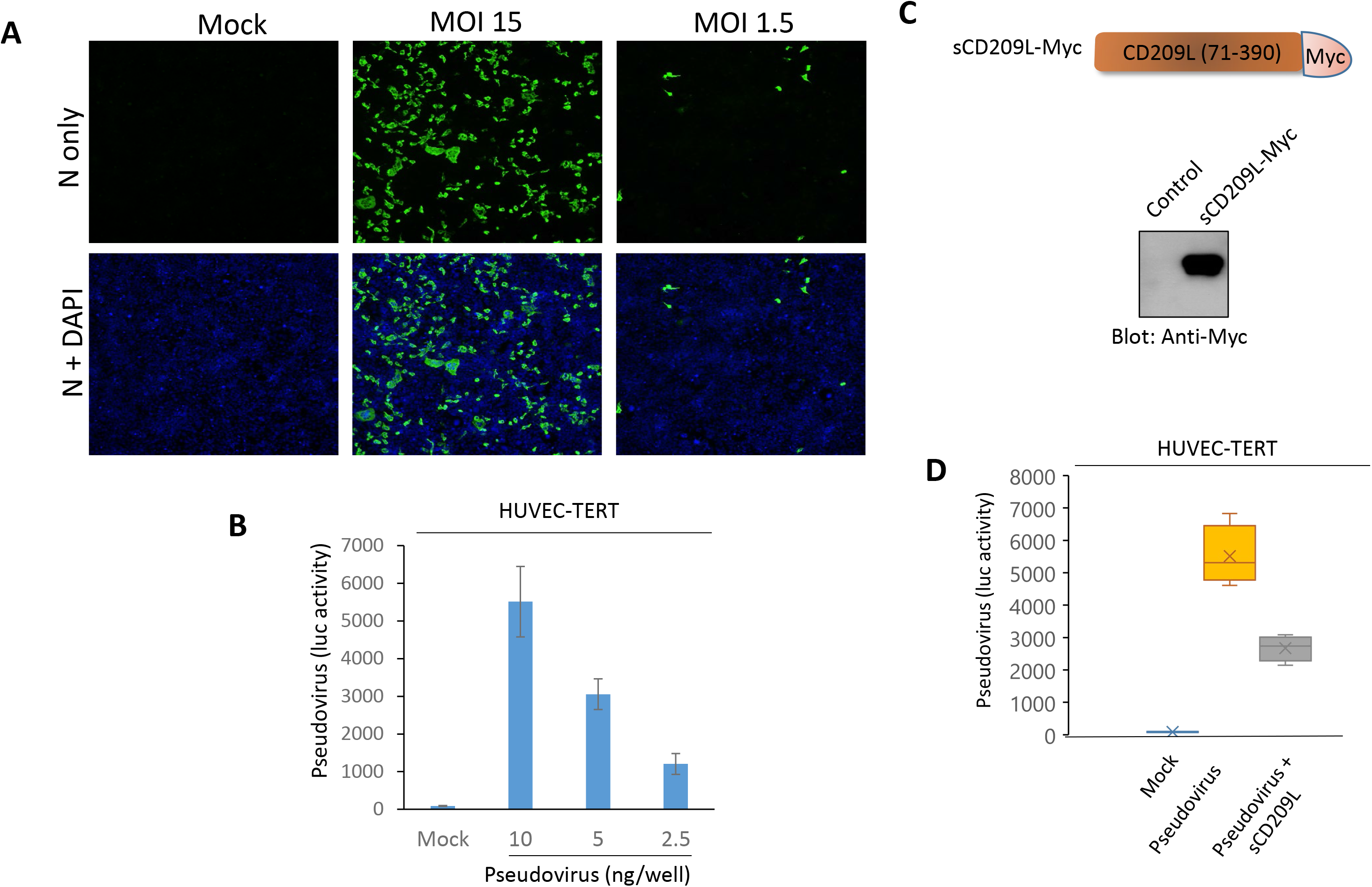
Endothelial cells are permissive to SARS-CoV-2 virus and soluble CD209L neutralizes viral entry. (**A**) HUVEC-TERT cells seeded in chamber slides were mock-infected or infected with SARS-CoV-2 at the indicated MOIs. Cells were fixed at one day post infection, and stained with an antibody directed against the viral nucleoprotein, N (green). Cell nuclei were stained with DAPI (blue). (**B**) HUVEC-TERT cells (2× 10^4^/well, 96-well plate, and quadruple/group) were infected with pseudovirus with different concentrations. After 24 h, cells were processed and subjected to luciferase activity assay and representative data are shown. (**C**) Schematic of Myc tagged soluble CD209 (sCD209L-Myc) and Western blot analysis of sCD209L-Myc. (**D**) HUVEC-TERT cells (2 × 10^4^/well, 96-well plate, and triplicate/group) were infected with mock, pseudotyped virus (10 ng/ml) with control conditioned medium (CM) or CM containing sCD209L (approximate concentration 1.5μg/ml). After 24 h, cells were analyzed for luciferase activity. #, P<0.05.

### CD209L mediates SARS-CoV-2 infection in endothelial cells

It has been proposed that the vascular system might be a direct target of SARS-CoV-2 infection ^8^. To explore if human endothelial cells are permissive to SARS-CoV-2, we infected HUVEC-TERT cells with SARS-CoV-2 at various multiplicities of infection (MOIs). A robust infection was observed at one day post infection, when the cells were infected with SARS-CoV-2 at a high MOI (MOI = 15). Even a lower MOI of 1.5 led to detectable infection levels at one day post infection, indicating that HUVEC-TERT cells are permissive to SARS-CoV-2 infection (**Figure 2A**). Moreover, we generated SARS-CoV-2 S-pseudotyped lentiviral particles and measured viral entry into HUVEC-TERT cells. SARS-CoV-2 S-pseudotyped lentiviral particles infected HUVEC-TERT cells in a concentration-dependent manner (**Figure 2B**). To investigate if CD209L expression in HUVEC-TERT cells promotes SARS-CoV-2 S-mediated entry, we carried out neutralization assays using a soluble form of CD209L (sCD209L (**Figure 2C**). sCD209L reduced viral entry by 48% (**Figure 2D**). Our data demonstrate that HUVEC-TERT endothelial cells are permissive to SARS-CoV-2 and sCD209L reduces SARS-CoV-2 S-pseudotyped viral entry.

To further examine the role of CD209L in viral entry, we infected HUVEC-TERT cells in which CD209L had been knocked down by shRNA and parental control cells with SARS-CoV-2 S-pseudotyped lentiviral particles. The result showed that loss of CD209L in HUVEC-TERT cells markedly reduced viral entry (**Figure 3A**). These data were confirmed by SARS-CoV-2 infection studies. Immunofluorescence analysis using an antibody directed against the viral nucleocapsid (N) protein revealed that silencing of CD209L led to a substantial decrease of SARS-CoV-2 infection in HUVEC-TERT cells (**Figure 3B**). Next, we asked whether expression of ACE2 in HUVEC-TERT cells contributes to viral entry by CD209L, albeit its expression in HUVEC-TERT cells is very low (**S. Figure 4A**). To address the relative contribution of ACE2 in SARS-CoV-2 entry in HUVEC-TERT cells, we knocked down ACE2 alone (**S. Figure 7A**) or CD209L alone by shRNA (**S. Figure 1B**) and subjected the cells to viral entry assay. The result showed that, while depletion of CD209L significantly reduced viral entry, knockdown of ACE2 had only a minor effect on the viral entry (**S. Figure 7B**), indicating that viral entry in HUVEC-TERT cells is mainly mediated by CD209L.

**Figure 3:**
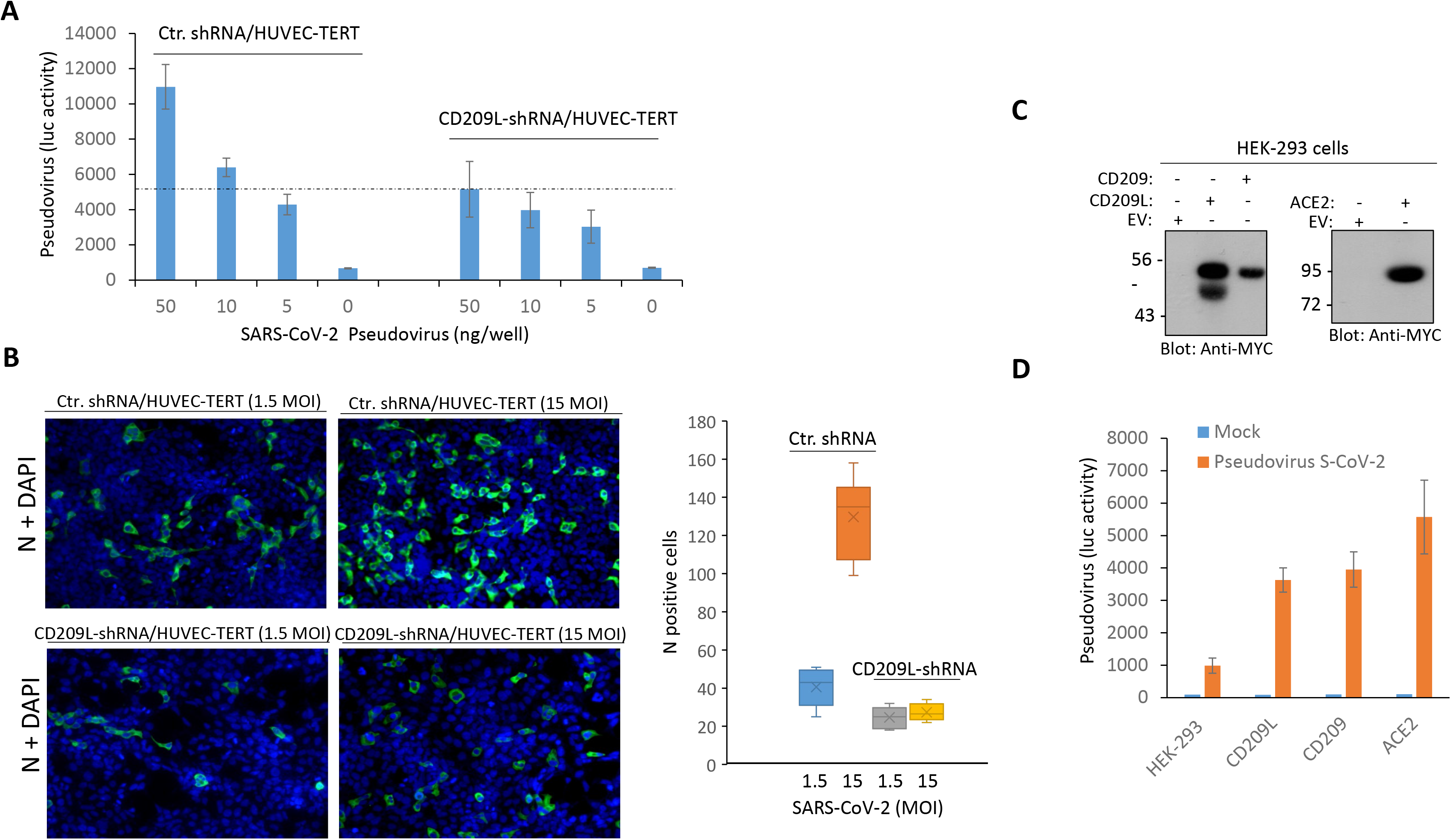
CD209L mediates SARS-CoV-2 entry and infection in endothelial cells. (**A**) HUVEC-TERT cells expressing control shRNA or CD209L-shRNA (2 × 10^4^/well, 96-well plate, and triplicate/group) were infected with different amounts of SARS-CoV-2 pseudotyped lentivirus. After 24 h, cells were processed and subjected to luciferase activity and representative data are shown. (**B**) HUVEC-TERT cells expressing control shRNA or CD209L-shRNA seeded in chamber slides (triplicate/group) were infected with SARS-CoV-2 at the indicated MOIs. Cells were fixed at one day post infection, and stained with an antibody directed against the viral nucleoprotein, N (green). Cell nuclei were stained with DAPI (blue). Quantification of N protein positive cells is shown. P<0.05. (**C**) Western blot analysis showing expression of CD209L, CD209 and ACE2. (**D**) HEK-293 cells expressing CD209L, CD209 or ACE2 (2 × 10^4^/well, 96-well plated, and quadruple wells/group) were infected with pseudotyped virus (10 ng/ml) or mock virus. After 24 h, cells were analyzed for luciferase activity. #, P<0.05.

To further illustrate the role of CD209L and CD209 in viral entry compared to ACE2, we over-expressed CD209L, CD209 or ACE2 in HEK-293 cells (**Figure 3C**) and compared the individual roles of these receptors in SARS-CoV-2 S mediated entry, using SARS-CoV-2 S-pseudotyped lentiviral particles. The results showed that both CD209L and CD209 were able to facilitate SARS-CoV-2 S-pseudotyped virus entry (**Figure 3D**). However, we consistently observed a higher S-pseudotyped virus entry in HEK-293 cells expressing ACE2 (**Figure 3D**). Altogether, our data demonstrate that CD2209L and CD209 can mediate SARS-CoV-2 entry, and thus even tissues lacking ACE2 can serve as infection sites.

### CD209L and CD209 act as receptors for SARS-CoV-2 spike

Given that CD209L facilitated SARS-CoV-2 entry, we investigated whether SARS-CoV-2 can physically interact with CD209L. Specifically, we asked whether the RBD domain of SARS-CoV-2-S binds to CD209L. We used multiple binding assays to determine if SARS-CoV-2 S interacts with CD209L. First, we generated a chimeric soluble S-RBD-Fc-Myc (**Figure 4A**) and tested its binding with CD209L in an immunoprecipitation assay using whole cell lysate derived from HEK-293 cells ectopically expressing CD209L (**Figure 4A**). The result showed that S-RBD-Fc domain binds to CD209L (**Figure 4B**). An unrelated Fc-chimeric protein, Fc-TMIGD1, did not bind to CD209L (**S. Figure 8A**), indicating that the binding of S-RBD-Fc-Myc with CD209L, is mediated by the S-RBD not by the Fc protein. To exclude that the observed binding between CD209L and S-RBD is a potential artifact of the Fc chimera system, we created a purified recombinant HIS tag S-RBD protein (**Figure 4C**) and analyzed CD209L binding via Far-Western blot analysis. The result showed that S-RBD-HIS strongly interacted with CD209L (**Figure 4D**). To further demonstrate the interaction of CD209L with S protein, we generated a S1-Myc construct which is composed of the N-terminal domain (NTB) and RBD (amino acids 19-685) and analyzed its binding with CD209L expressed in HEK-293 cells. The result showed that, similar to Fc-S-RBD, S1 binds to CD209L in a similar manner, indicating that RBD is the main domain involved in the binding of S protein with CD209L (**S. Figure 9B**). Additionally, S-RBD interacted with CD209L endogenously expressed in HUVEC-TERT cells (**S. Figure 9C**), indicating that the binding of S-RBD with CD209Lexpressed in HEK-293 cells is not an artifact of over-expression of CD209L in HEK-293 cells. Next, we compared the binding of S-RBD-HIS with CD209L, CD209 and ACE2 in a dot blot assay. S-RBD-HIS interacted with CD209L, CD209 and ACE2 in a concentration-dependent manner (**Figure 4E**). The strongest signals were observed with Fc-ACE2-Myc (**Figure 4E**), suggesting a higher affinity between S-RBD and ACE2 compared to CD209L and CD209.

**Figure 4:**
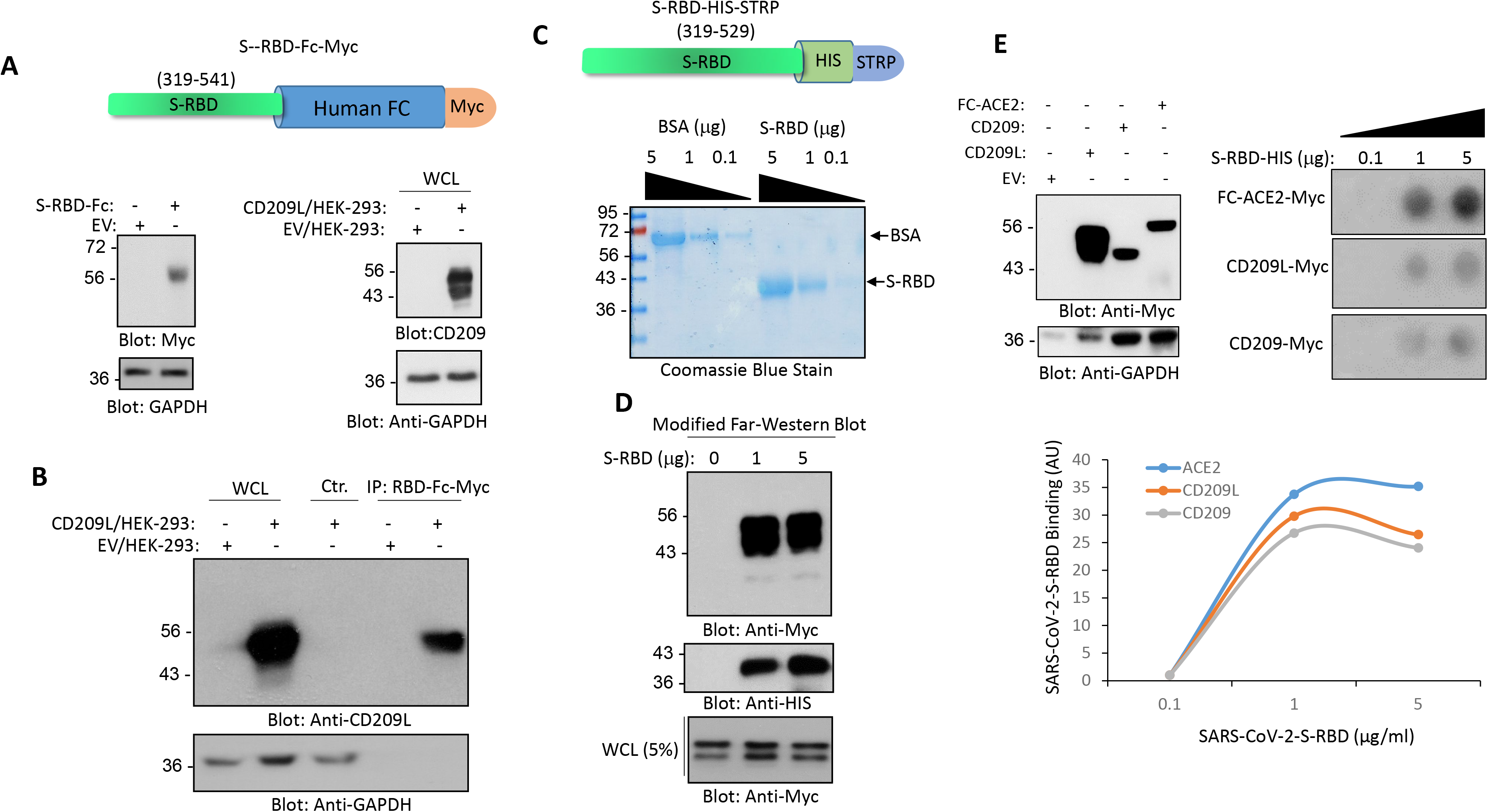
CD209L and CD209 bind to SARS-CoV-2-S-RBD. (**A**) Schematic of Fc-CoV-2-S-RBD, expression of Fc-CoV-2-S-RBD and CD209L in HEK-293 are shown. (**B**) Immunoprecipitation assay demonstrates the binding of CD209L with Fc-CoV-2-S-RBD. (**C**) Schematic and Coomassie blue stain of CoV-2-S-RBD-HIS are shown. (**D**) Far-Western blot analysis shows the binding of HIS-STRP tagged CoV-2-S-RBD with CD209L. (**E**) Western blot analysis of CD209L, CD209 and Fc-ACE2 ectopically expressed in HEK-293 cells. (**F**) CoV-2-S-RBD-HIS applied onto PFVD membrane with varying concentrations as indicated in the figure legend via Dot blot apparatus. The membranes, after blocking with 5% BSA, were incubated with cell lysates derived from HEK-293 cells expressing Fc-ACE-2-Myc, CD209L-Myc or CD209-Myc and the binding of ACE2, CD209L and CD209 to CoV-2-S-RBD was detected with anti-Myc antibody. Quantification of the dot blots is shown. AU, arbitrary unit. Image J software was used to quantify the dot blots.

ACE2 is described as a main entry receptor for SARS-CoV-2 ^13^; we asked whether CD209L can associate with ACE2. To answer this question, we carried out a co-immunoprecipitation assay and demonstrated that CD209L can form a heterodimer with ACE2 (**S. Figure 10**), suggesting both ACE2-independent and ACE2-dependent roles for CD209L in SARS-CoV-2 entry and infection. However, whether CD209L directly interacts with ACE2 or its interaction with ACE is aided by additional proteins requires further investigation. To investigate the mechanism of CD209L interaction with ACE2, we asked whether the CRD domain on CD209L is involved in the binding of ACE2. We treated HEK-293 cells expressing CD209L-Myc with EGTA that specifically removes calcium and, as a result, disables the mannose recognition capability of the CRD domain. The result showed that the removal of calcium had no significant impact on the binding of ACE2 with CD209L, indicating that the CRD is not involved in the CD209L interaction with ACE2 (**S. Figure 11**).

### CD209L is *N*-glycosylated and *N*-glycosylation interferes with interaction of CD209L with SARS-CoV2 Spike Protein

CD209L interacts with glycoproteins via its C-type lectin domain and it is itself subject to *N*-glycosylation^37^. We determined the occupancy of the potential *N*-glycosylation sites on our CD209L construct and investigated whether *N*-glycosylation plays a role in the interaction of CD209L with spike protein. After treatment of the immunoprecipitated protein with PNGase F in the presence of H_2_^18^O (which removes *N*-linked glycans and converts N→D, incorporating ^18^O at the formerly-glycosylated site, resulting in a mass shift of 3 u), we detected only the formerly-glycosylated version of the peptide spanning the N92 *N*-glycosylation sequon (**S. Figure 12**). The results indicated that the CD209L protein is fully *N*-glycosylated at site N92. Furthermore, nUPLC-MS/MS analyses of CD209L digests enabled detection of a glycopeptide bearing high-mannose type *N*-linked glycosylation at site N92 (**Figure 5A**). The dominant glycoform was Man8. In contrast, although the peptide that includes the N361 site was observed, no deamidation/^18^O incorporation was detected after the PNGase F treatment. (**S. Figure 12**). This result indicated that the N361 site is unoccupied.

**Figure 5:**
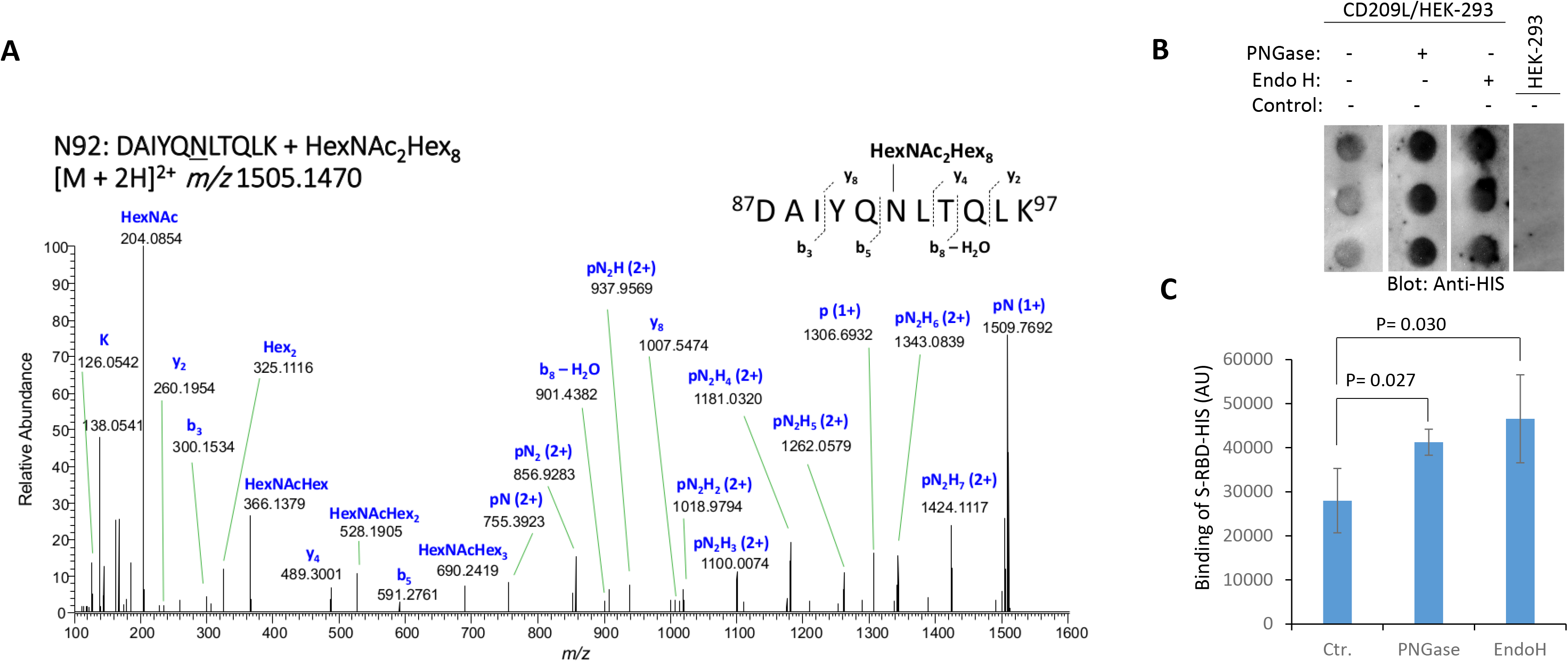
CD209L *is N-glycosylated and N*-glycosylation modulates its interaction with SARS-CoV2 Spike Protein. **(A)** HCD MS/MS spectrum of *m/z* 1505.1470, assigned as the [M + 2H]^2+^ of the glycopeptide ^87^DAIYQNLTQLK^97^ + HexNAc_2_Hex_8_, corresponding to CD209L site N92 bearing a Man_8_ glycan. Stepped-collision energy (15/25/35% NCE) was used. The fragment ions do not retain the glycan moiety. p = peptide, N = *N*-acetylhexosamine (HexNAc), H = hexose (Hex). **(B)** Dot blot of CD209L/HEK-293 cell lysate under control conditions (no treatment), or treated with PNGase F or Endo H. Dot blot of HEK-293 cell lysate was used as a negative control. Following treatment of lysate, the spike protein receptor-binding domain (S-RBD-HIS) was incubated with the immobilized lysate, followed by detection with anti-HIS antibody. **(C)** Quantification of blots is shown. AU, arbitrary unit. Image J software was used to quantify the dot blots

To investigate the role of CD209L *N*-glycosylation in its interaction with the SARS-CoV-2-S-RBD, a dot blot analysis was performed in which CD209L/HEK-293 or control HEK-293 cell lysates were immobilized on a PVDF membrane, followed by incubation with the spike protein receptor-binding domain (S-RBD-HIS). Treatment with PNGase F or Endo H, which remove both complex and high-mannose *N*-linked glycans or only high-mannose *N*-linked glycans, respectively, led to an increase in the binding of SARS-CoV-2 S-RBD in a statistically significant manner compared to control conditions (**Figure 5B, C**). Together, the data suggest that high-mannose *N*-linked glycans on CD209L and/or the SARS-CoV2 S-protein hinder the interaction of CD209L and the S-protein, while the absence of high mannose glycans favors the CD209L-S protein interaction.

## Discussion

In this study, we demonstrate that CD209L is broadly expressed in human pulmonary and kidney tissues, in both endothelial cells and epithelial cells. CD209L and CD209 can serve as receptors for SARS-CoV-2, as evidenced by their interactions with the purified RBD domain of the SARS-CoV-2 S. Recognition of CD209L and CD209 by the RBD domain of SARS-CoV-2 S mediates virus entry. Endothelial cells that lack ACE2 are permissive to SARS-CoV-2 infection. Interfering with CD209L expression or function with shRNA or soluble CD209L in endothelial cells inhibited SARS-CoV-2 virus entry. Similarly, ectopic expression of CD209L and CD209 in HEK-293 cells increased SARS-CoV-2 entry, which underscores the potential roles of these receptors in SARS-CoV-2 in infection of human cells. This idea is consistent with the previous studies which demonstrated that CD209L interacts with Ebola virus surface glycoprotein and mediates endothelial cell infection ^45–46^. The actions of host viral entry receptors, through multiple mechanisms such as endocytosis and fusion, can result in viral replication in host cells, whereas cell surface structures (*e.g*., glycans and lectins) can serve as adhesion receptors binding to virus and significantly enhancing virus entry into the target cells *via* their interactions with features on the viral surface (*e.g.*, spike protein) or with the entry receptors (*e*.*g*., ACE2). CD209L and CD209 have been described as attachment factors for multiple viruses, including Marburg and SARS ^25, 47^. Subsequent studies demonstrated CD209L and CD209 themselves can act as entry receptors for SARS independent of ACE2 ^32, 48^. Our results indicate that CD209L and CD209, by acting as entry receptors, mediate trans-infection of SARS-CoV-2 in endothelial cells. However, CD209L in other cell types could function as an attachment factor, but the present study have not investigated this mechanism of CD209L in viral entry. Interestingly, a recent study demonstrates that CD209L can also mediate cis-infection of SARS-CoV-2 in monocytes and T-lymphocytes ^49^. Another important aspect of CD209L involvement in SARS-CoV-2 infection is the demonstration of the interaction of CD209L with ACE2. This suggests that CD209L is capable of facilitating SARS-CoV-2 entry in both ACE2-dependent and - independent manners. However, further detailed studies are needed to fully elaborate the ACE2-independent function of the CD209L and CD209 in SARS-CoV-2 viral entry and infection.

When compared to CD209L or CD209, ACE2 appeared to have a stronger binding to SARS-CoV-2 and also mediated SARS-CoV-2 entry in HEK-293 cells more efficiently. Nevertheless, in agreement with the proposed role of CD209L and CD209 in SARS-CoV-2 infection, multiple biochemical assays demonstrated the direct physical interaction of CD209L and CD209 with SARS-COV-2 S-RBD protein. CD209L and CD209 showed similar interactions with SARS-CoV-2 S-RBD. Thus, CD209L and CD209 can serve as alternative entry receptors for SARS-CoV-2 spike protein. In agreement with our findings, a recent report by van Kooyk and colleagues also reported an ACE2-independent binding between CD209L and SARS-CoV-2 S protein^50^. Furthermore, while the preprint of this manuscript had been shared in the bioRxiv (https://www.biorxiv.org/content/10.1101/2020.06.22.165803v1) another group also reported the binding of SARS-COV-2 S protein with CD209 in a glycosylation-dependent manner, although they did not determine the site occupancies or the glycoform distributions^51^. Another important feature of CD209L interaction with SARS-CoV-2 S protein is that *N*-glycosylation of CD209L appears to hinder its interaction with the SARS-CoV-2 S protein, suggesting that differential *N*-glycosylation of CD209L (and/or the S-protein) due to differences in cell types, pathology, individuals or species could influence the overall interaction of CD209L with SARS-CoV-2. One possible mechanism for the function of *N*-glycosylation of CD209L is that *N*-glycosylation generates a hindrance for the CRD domain and hence reduces its ability to recognize the mannose-rich SARS-CoV-2 S protein. However, further structure-function studies are needed to elucidate the possible mechanism by which *N*-glycosylation of CD209L impacts its recognition of SARS-CoV-2-S protein.

ACE2 was originally reported to be expressed in various cell types, including endothelial cells of the heart, kidneys, and testis ^52^ and in lung alveolar epithelial cells^14^. However, more recent studies showed that ACE2 is expressed relatively at low levels or is undetectable in endothelial cells of lung, liver, skin and intestine^16^, and lung tissue ^15^. We observed a strong staining of CD209L in alveolar type II epithelial lung cells, renal proximal epithelial cells and blood vessels, which might provide potential routes of entry for SARS-CoV-2. However, we did not observe CD209 expression in blood vessels or lung epithelial cells. Previous studies have shown that CD209 is mostly expressed in dendritic cells, tissue-resident macrophages, and B cells ^53–54^, suggesting that these cell types could be targeted by SARS-CoV-2 via recognition of CD209.

Recent studies suggest that the vascular system is a major site of attack by SARS-CoV-2 ^7–8^. COVID-19 patients suffer from distinct endothelial cell injury (*i.e*., endothelitis), and altered angiogenesis ^10^ with widespread microvascular thrombosis ^3–4^. These observations, coupled with the fact that vascular endothelial dysfunction also plays crucial roles in the pathogenesis of COVID-19 ^55^, underscores the role of endothelial cells in the pathobiology of SARS-CoV-2 infection and also a therapeutic opportunity.

## Conclusion

Our present study reveals that CD209L not only can act as a SARS-COV-2 entry receptor, but also performs critical functions in the angiogenic responses of endothelial cells. This suggests that SARS-COV-2, by exploiting CD209L, could subvert CD209L function in endothelial cells, leading to endothelial cell injury and altered angiogenesis. However further studies such as cellular context and physical interaction of CD209L and CD209 with spike protein and ACE2 are required to fully understand the role of CD209 and CD209L in SARS-CoV-2 as we used only a cell line to link CD209L to viral entry in endothelial cells. Investigations of the molecular and cellular effects of SARS-CoV-2 on the vascular system are thus critical to defining the selective roles of CD209L and CD209, in both the presence and absence of ACE2 in the this and other systems, in order to explore the therapeutic potential of CD209L and CD209 as targets for combatting COVID-19.

## Materials and Methods

Full details of the plasmids, antibodies and procedures used are provided in the supplemental data. The mass spectrometry proteomics data have been deposited to the ProteomeXchange Consortium via the PRIDE ^56^ partner repository with the dataset identifier PXD021309.

### Statistical analyses

Experimental data were subjected to Student t-test or One-way analysis of variance analysis, where appropriate, with representation from at least three independent experiments. p<0.05 was considered significant.

## Supporting Information

Twelve additional figures including material and methods.

## Safety Statement

No unexpected or unusually high safety hazards were encountered.

## Acknowledgement

This work was supported in part through grants from the National Institutes of Health (NIH) National Cancer Institute R21 CA191970 and R21 CA193958, CTSI grant 1 UL1 TR001430 and Malory Fund, Department of Pathology, Boston University (to N.R.), a grant from The Evans Center for Interdisciplinary Biomedical Research ARC on COVID-19, and NIH grants from the National Institute for Allergy and Infectious Diseases (NIAID) R01 AI064099 and National Institute for Aging R01 AG060890 (to S.G.), NIH grants from NIAID R01 AI133486 and National Center for (NCATS) CTSI UL1 TR001430, and awards from Evergrande MassCPR sub-award 280870.5116795.0025 and Fast Grants (to E.M.), NIH Heart, Lung and Blood Institute grant R01 HL132325 (to V.C.C.), NIH grants from the Institute of General Medical Sciences R24 GM134210, the Office of the Director S10 OD021728 and BUSM CTSI 1 UL1 TR001430 COVID-19 Related Research Award (to C.E.C.), NIH/NIAID grant R01 AI146779 and a Massachusetts Consortium on Pathogenesis Readiness (MassCPR) grant (to A.G.S.) and NIH training grants: T32 GM007753 for B.M.H. and T.M.C; T32 AI007245 for J.F.; and T32 HL007035 for E.L.S. The content is solely the responsibility of the authors and does not necessarily represent the official views of the funding agencies.

## Authors’ contributions

NR, RA, EM, SG, VCC, KBC and CEC were involved in the experimental design, data interpretation, and writing and editing of the manuscript. NR, RA, MAN, WY, JB, ES, QZ, JO, KBC and CX performed the experiments. JF, BMH, TMC and AGS provided protein and reagents.

## Data availability

Data and reagents are available from the corresponding author upon request. Mass spectrometry data are available via ProteomeXchange with identifier PXD021309.

## Synopsis

Understanding the interactions between SARS-CoV-2 with host cells is of high importance. ACE2 is recognized as a major entry receptor, but SARS-CoV-2 may also employ alternative receptors for cell entry and these may hold the key to infection in tissues where ACE2 has a low expression level or is absent. We identify CD209L/L-SIGN and CD209/DC-SIGN as receptors for SARS-CoV-2. We show that CD209L is *N*-glycosylated and this modification modulates the binding of CD209L with spike protein. CD209L interacts with ACE2, suggesting that CD209L and ACE2 could function as co-receptors for SARS-CoV-2 entry and infection. Human endothelial cells are permissive to SARS-CoV-2 infection. We show that interfering with CD209L activity in endothelial cells by knockdown or with introduction of soluble CD209L inhibits virus entry, suggesting a novel target for development of antiviral drugs.

## Graphical ABSTRACT

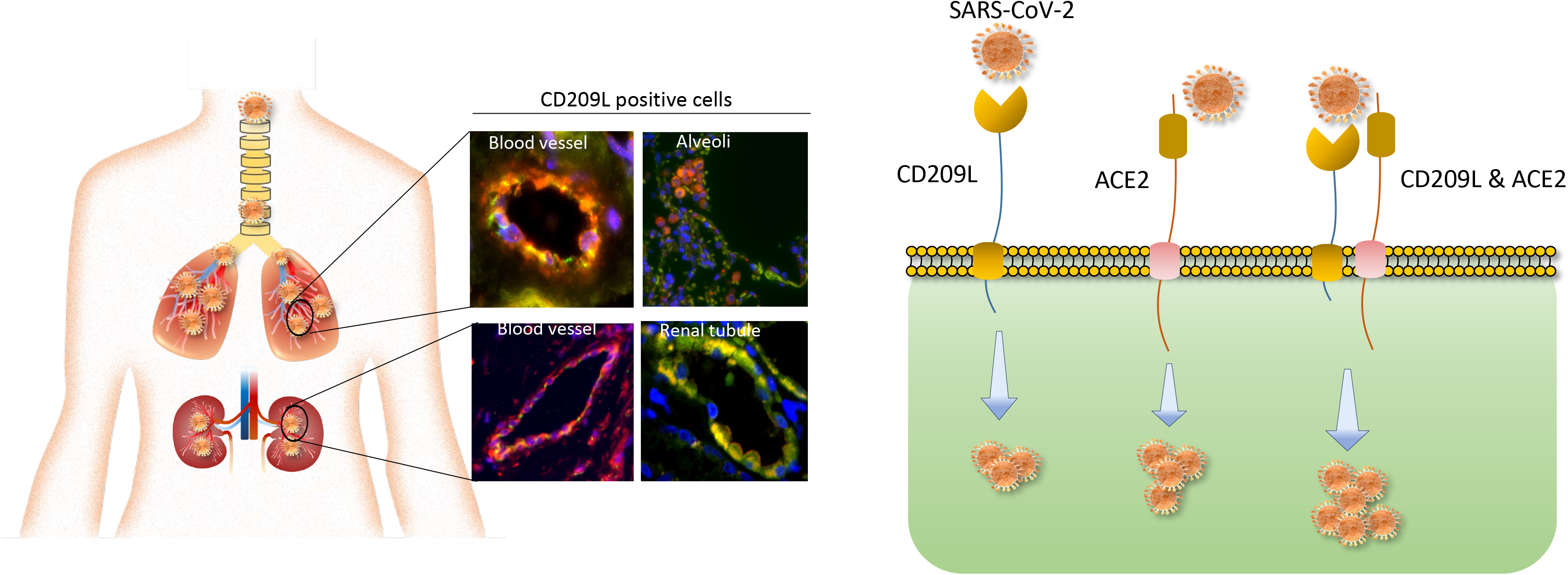

