## Supplemental data for "CD209L/L-SIGN and CD209/DC-SIGN act as receptors for SARS-CoV-2"

[1]Department of Pathology, School of Medicine, Boston University Medical Campus, Boston, MA 02118. [2]Renal Section, Department of Medicine, Boston University Medical Center, Boston, MA. [3]Department of Microbiology, Boston University School of Medicine, Boston, MA. [4]National Emerging Infectious Diseases Laboratories (NEIDL), Boston University, Boston, MA. [5] Center for Biomedical Mass Spectrometry, Boston University School of Medicine, Boston, MA 02118. [6]Ragon Institute of MGH, MIT, and Harvard, Cambridge, MA 02139. [7]Department of Microbiology, Harvard Medical School, Boston, MA 02115.

**Short Title**: CD209L and CD209 are receptors for SARS-CoV-2

***Corresponding authors:**

Prof. Catherine E. Costello
Center for Biomedical Mass Spectrometry
Boston University School of Medicine
670 Albany St, rm 511
Boston, MA 02118-2646


Prof. Nader Rahimi
Department of Pathology
Boston University School of Medicine
670 Albany St., rm 510
Boston, MA 02118-2646


**S. Figure 1**. Staining of human lung tissue with control antibody and validation of CD209L antibody Page 2
**S. Figure 2**. CD209L is expressed in human renal proximal epithelial cells Page 3
**S. Figure 3**: CD209 expression in lung and renal tissues Page 4
**S. Figure 4**. CD209L is expressed in human endothelial cells Page 5
**S. Figure 5**. CD209L regulates angiogenic responses of endothelial cells Page 6
**S. Figure 6**. CD209L regulates cytoplasmic protrusions of HUVEC-TERT cells Page 7
**S. Figure 7**. Effect of knockdown of ACE in SARS-CoV-2 in endothelial cells. Page 8
**S. Figure 8**. Unrelated Fc chimeric protein, TMIGD1, does not bind to CD209L Page 9
**S. Figure 9**. SARS-CoV-2 S1 and S-RBD-Fc-Myc bind to CD209L Page 10
**S. Figure 10**. CD209L interacts with ACE2 Page 11
**S. Figure 11**: Binding of CD209L with ACE2 does not require functional CRD domain of CD209L Page 12
**S. Figure 12**. MS/MS spectra of rCD209L peptides with potential *N*-linked glycosylation sites, after treatment with PNGase F/$H_2{}^{18}O$ Page 13
**Materials and methods**: Page 14

**S. Figure 1. Staining of human lung tissue with control antibody**. (**A**) No noticeable staining was observed with control antibody. (**B**) Western blot analysis of cell lysates from HUVEC-TERT cells transfected with control shRNA or CD209L-shRNA.

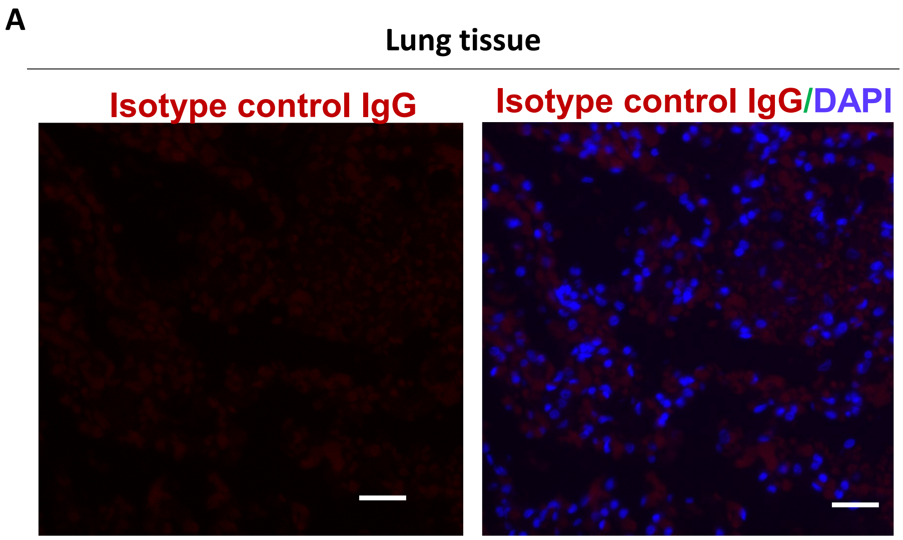


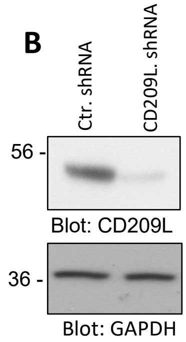

**S. Figure 2. CD209L is expressed in human renal proximal epithelial cells**. (**A**) Renal tissue stained with aquaporin1 (green) and CD209L (red). (**B**) Staining of CD209L in renal cortex, which is negative. Glomeruli (G). DAPI (Blue). L, lumen

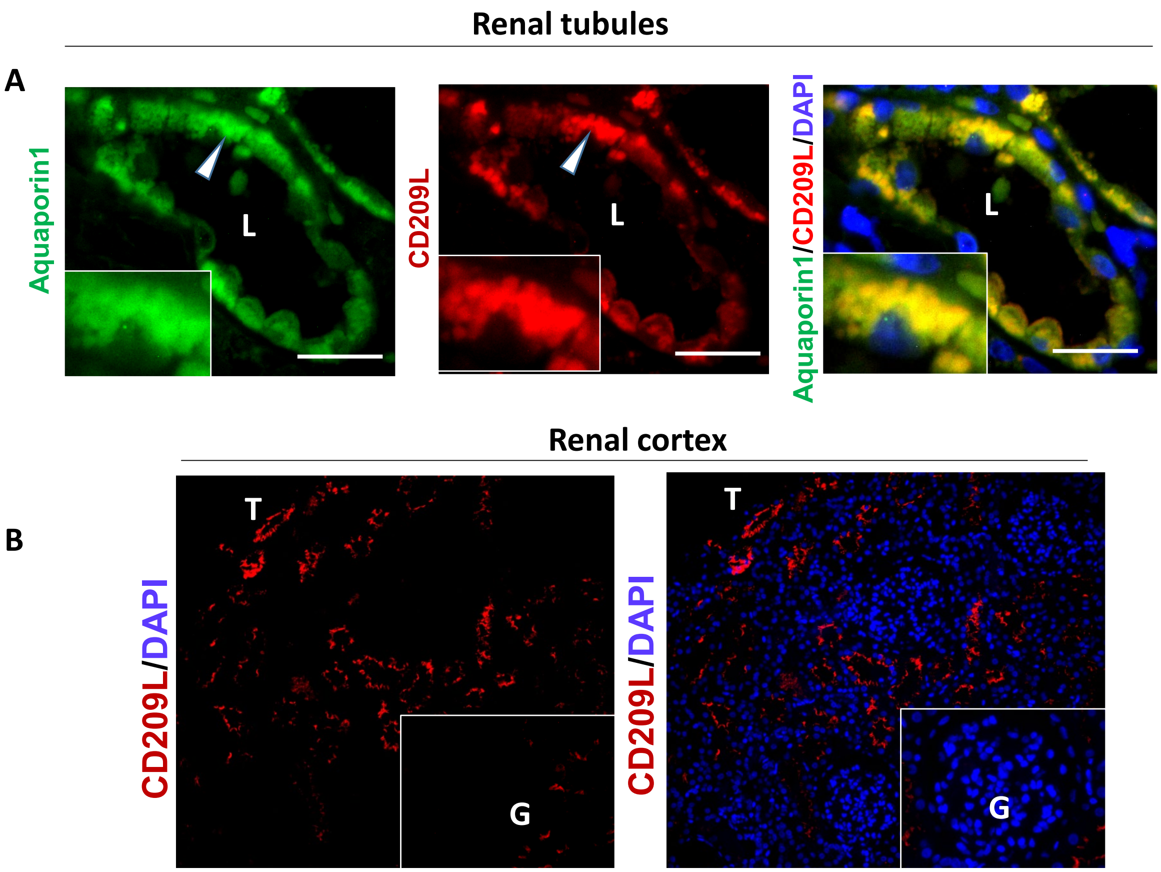

**S. Figure 3: CD209 expression in lung and renal tissues**: PFA fixed human lung and kidney tissues were subjected to immunofluorescence staining and stained with anti-MUC1, CD31, and CD209 antibodies. (**A**) Lung alveoli epithelial cells were mostly negative for CD209. (**B**) Staining of renal proximal epithelial cells with anti-aquaporin2 and anti-CD209 antibodies. (**C**) Glomerulus capillaries were mostly negative for CD209. Glomerulus (G), Lumen (L), Alveoli (A).

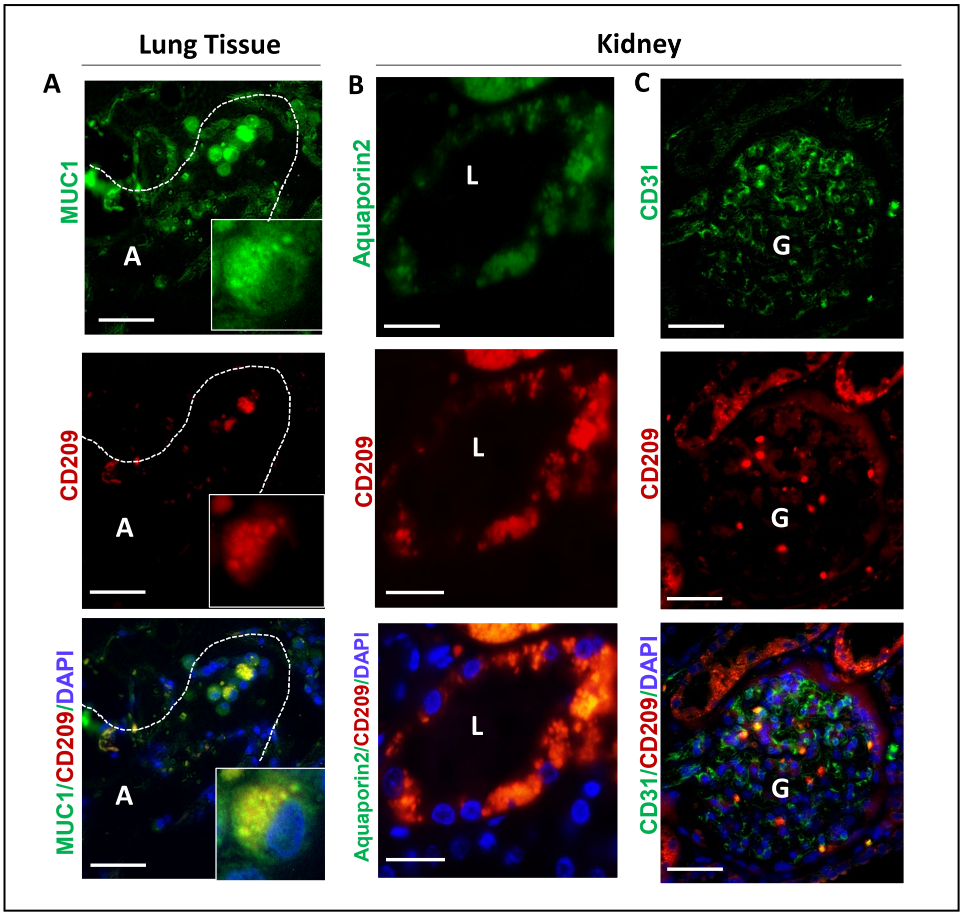

**S. Figure 4. CD209L is expressed in human endothelial cells**. (**A**) Western blot analysis of CD209L. CD209L detected in HUVEC-TERT cells but not in PAE or COS1 cells. NS, non-specific. (**B**) Western blot analysis of whole cell lysates derived from clung carcinomas A549, H1299 and HUVEC-TERT cells. (**C**) qPCR analysis of CD209L in HUVEC-TERT cells expressing control shRNA or CD209L-shRNA.

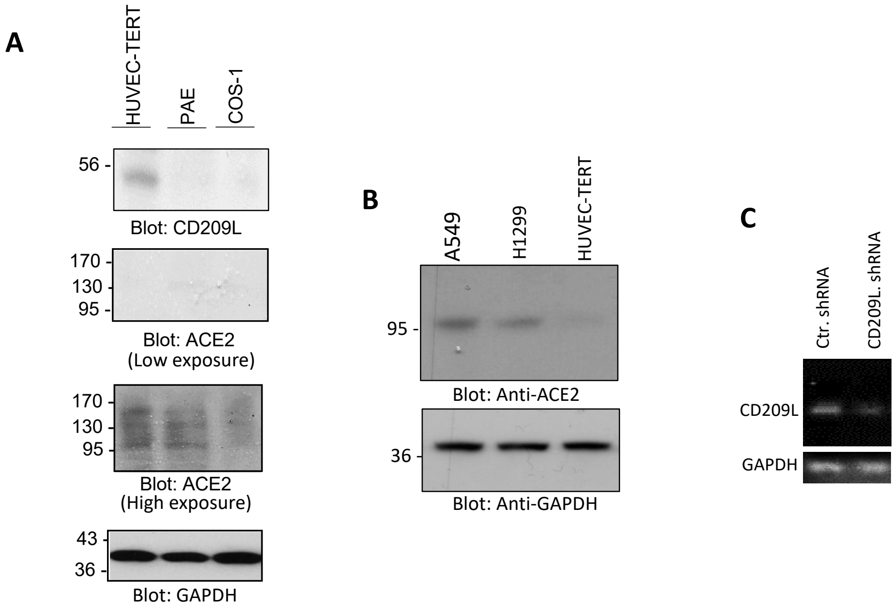

**S. Figure 5. CD209L regulates angiogenic responses of endothelial cells**. (**A**) HUVEC-TERT cells expressing control shRNA or CD209L-shRNA were seeded on 96-well plates (triplicate/group) coated with sDC209L. After 30 minutes, cells were fixed, stained crystalline blue and number of adherent cells were counted and representative graph is shown. (**B**) HUVEC-TERT cells expressing control shRNA or CD209L-shRNA were seeded on 96-well plates (triplicate/group) coated with Matrigel and capillary tube formation was analyzed after 6 hours. Image J used to quantify tube formation. (**C**) Fully confluent HUVEC-TERT cells expressing control shRNA or CD209L-shRNA seeded on 6-well plates (triplicate/group) were subjected to migration/wound healing assay and pictures were taken after 6 or 12 hours and representative images are shown.

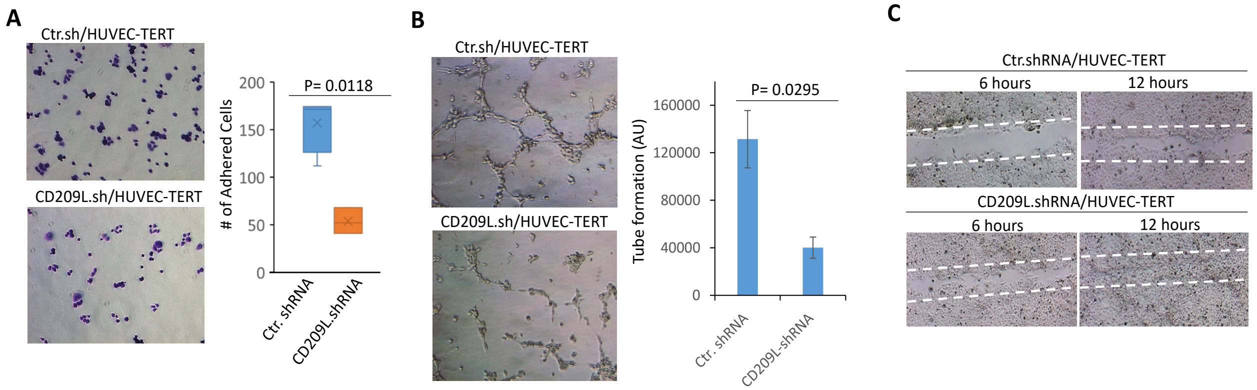

**S. Figure 6. Knockdown of CD209L alters cytoplasmic protrusions at the leading edge of HUVEC-TERT cells**. Phalloidin staining of HUVEC-TERT cells expressing control shRNA or CD209L-shRNA. The arrows show an increase in the cytoplasmic protrusions at the leading edge of HUVEC-TERT cells expressing CD209L-shRNA.

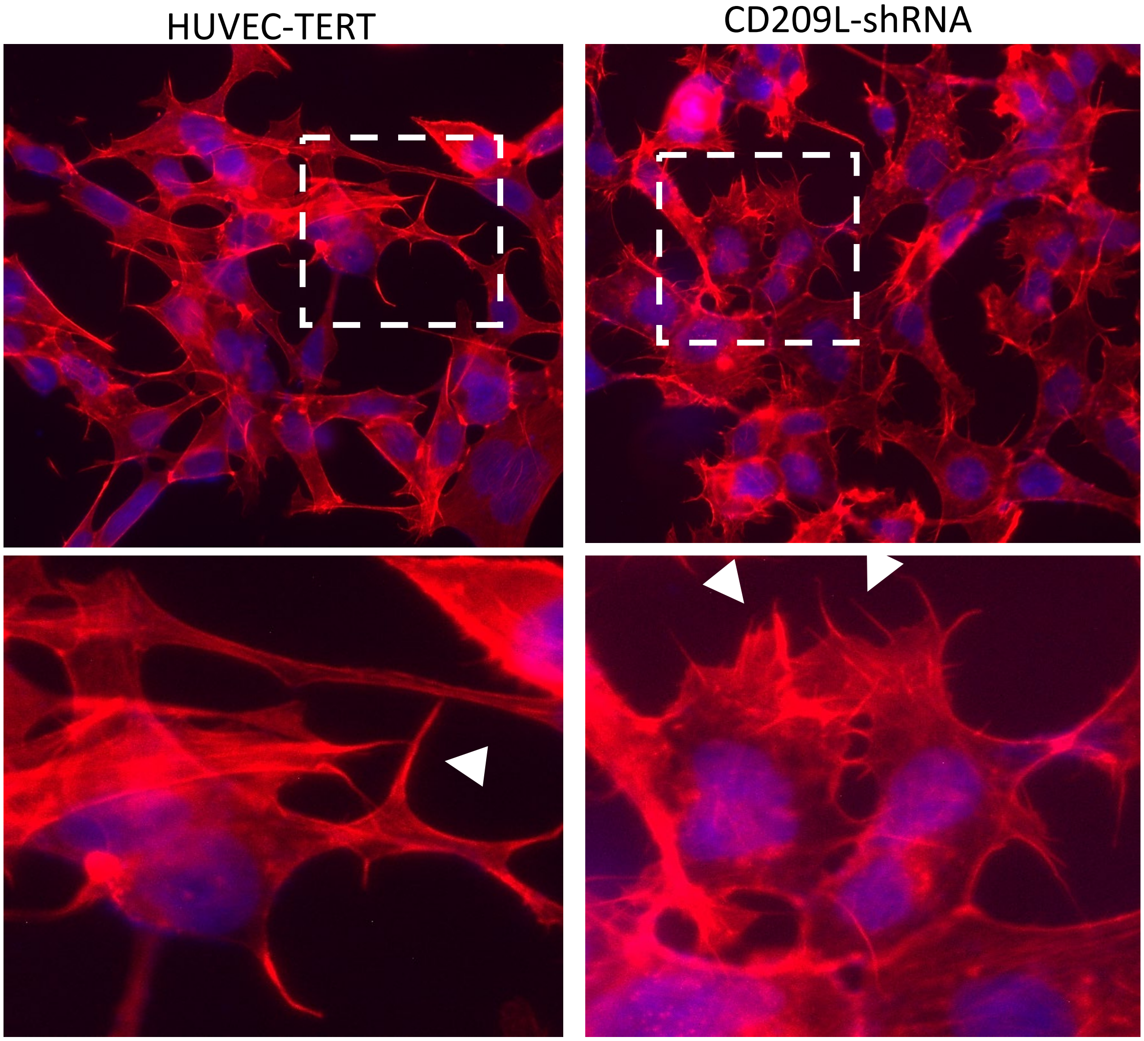

**S. Figure 7. Effect of knockdown of ACE in SARS-CoV-2 in endothelial cells**. (**A**) Western blot analysis of whole cell lysates derived from HUVEC-TERT cells transfected with control shRNA or ACE-2 shRNA. (**B**) HUVEC-TERT cells expressing control shRNA, CD209L-shRNA or ACE2-shRNA (2 X $10^4$/well, 96-well plate, and quadruple/group) were infected with different amounts of SARS-CoV-2 pseudotyped lentivirus. After 24 h, cells were processed and subjected to luciferase activity and representative data are shown.

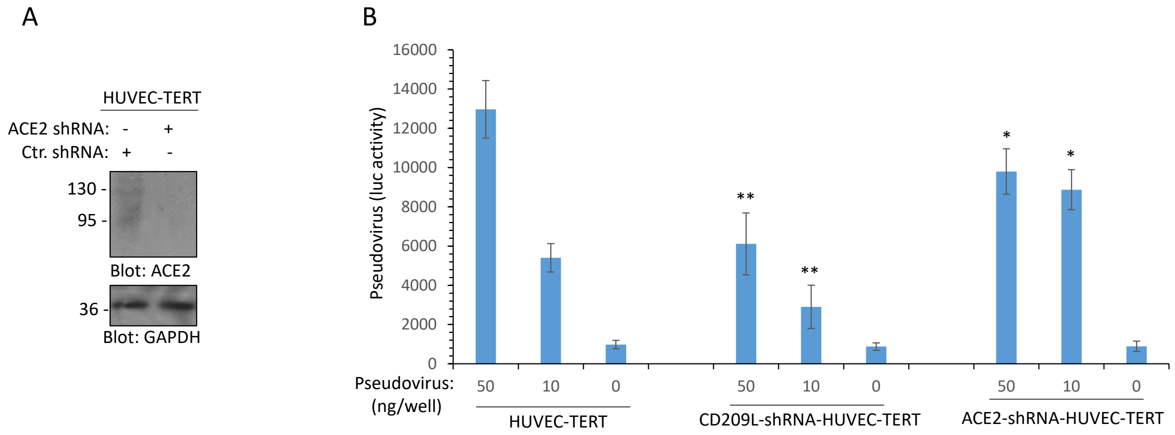

**S. Figure 8. Unrelated Fc chimeric protein, TMIGD1, does not bind to CD209L. (A)** Immunoprecipitation assay demonstrates that TMIGD1-Fc-Myc does not bind to CD209L.

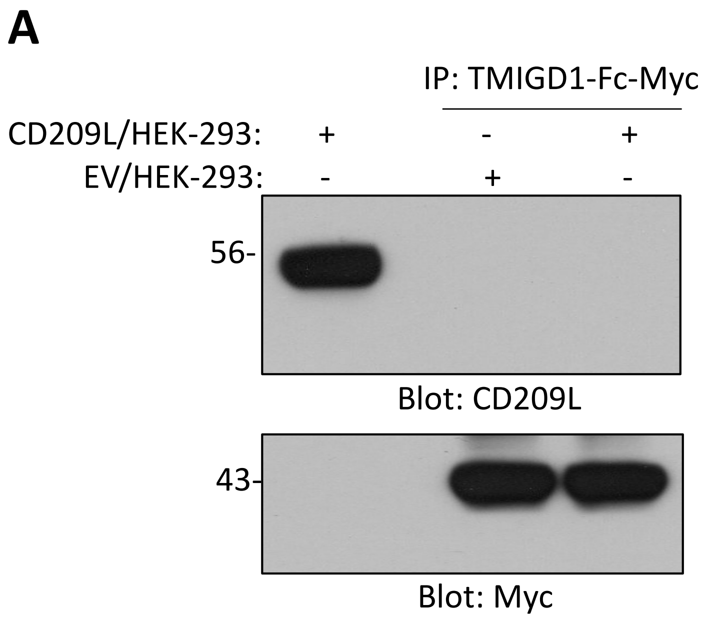

**S. Figure 9. SARS-CoV-2 S1 and S-RBD-Fc-Myc bind to CD209L: (A)** Schematic of construction of S-RBD-Myc and S1-Myc are shown. (**B**) Immunoprecipitation assay demonstrates that S-RBD-Fc-Myc and S1-Myc bind to CD209L in a similar manner. **(C)** Whole cell lysates (WCL) from HUVEC-TERT cells expressing control shRNA or CD209L-shRNA were subjected to immunoprecipitation assay using S-RBD-HIS followed by immunoblotting with anti-CD209L antibody. The same membrane was blotted for S-RBD-HIS and loading control, GAPDH.

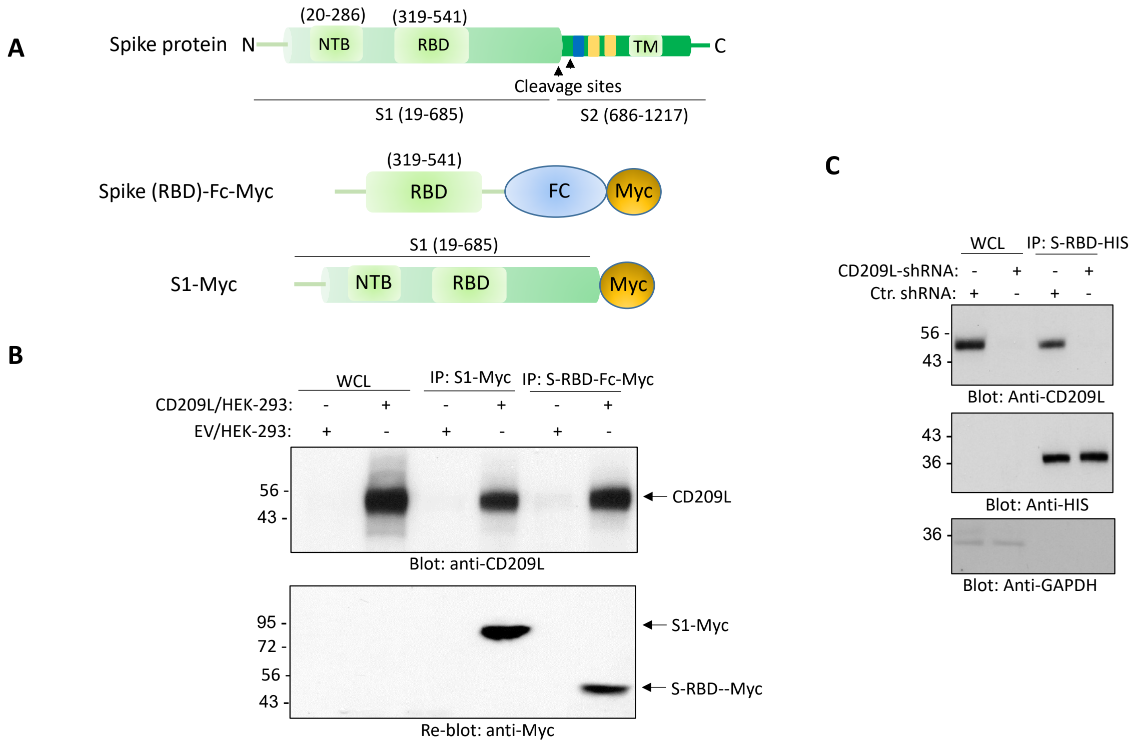

**S. Figure 10. CD209L interacts with ACE2**: Equal number of whole cell lysates derived from HEK-293 cells expressing CD209L-Myc or ACE2 or mixed (1:1 ratio) and subjected to co-immunoprecipitation using anti-c-Myc magnetic bead followed by Western blotting using anti-ACE2. The same membrane was also re-blotted with Anti-Myc for CD209L level. Whole cell lysates (WCL) used as a loading control and also blotted with GAPDH for protein loading control.

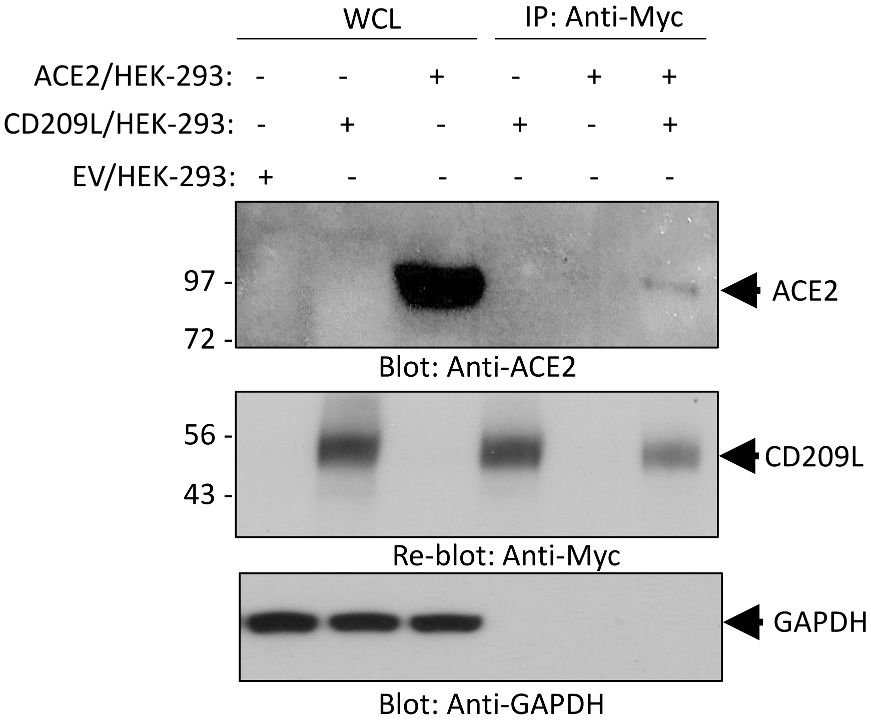

**S. Figure 11: Binding of CD209L with ACE2 does not require functional CRD domain of CD209L:**

HEK-293 cells expressing CD209L treated with calcium chelator, EGTA (5mM for 5 min). Cells were lysed and subjected to co-precipitation assay as described in S. Figure 10. EGTA treatment had only a negligible effect on the binding of CD209L with ACE2.

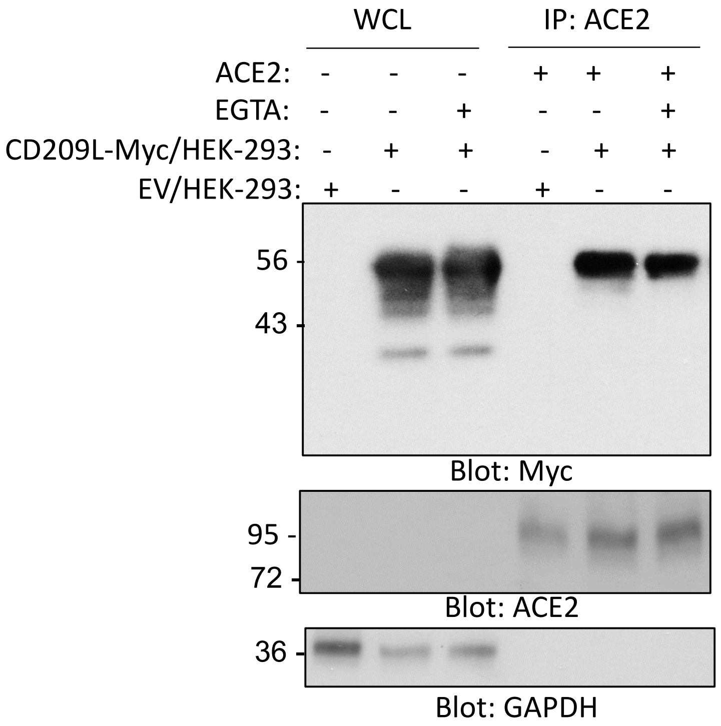

**S. Figure 12. MS/MS spectra of peptides containing the potential *N*-linked glycosylation sites from recombinant CD209L expressed in HEK-293 cells, after treatment with PNGase F/$H_2{}^{18}O$**, verify the peptide sequences and indicate occupancy. (a) HCD MS/MS spectrum of $[M + 3H]^{3+}$ *m/z* 890.0931 assigned to the tryptic peptide containing residues 75-97. The mass shift compared to the unmodified peptide VPSSLSQEQSEQDAIYQNLTQLK (+2.9883 u) corresponds to the conversion of N92→D92 with incorporation of $^{18}O$, the change expected for removal of an *N*-linked glycan in the presence of $H_2{}^{18}O$. (b) HCD MS/MS spectrum of $[M + 3H]^{3+}$ *m/z* 1021.7328 corresponds to the amino acid sequence $^{354}$YWNSGEPNNSGNEDC(CAM)AEFSGSGWNDNR$^{380}$, where Cam = carbamidomethyl, and shows no mass shift that would indicate N361→D361 conversion or $^{18}O$ incorporation, indicating there was no detectable elimination of an *N*-linked glycan.

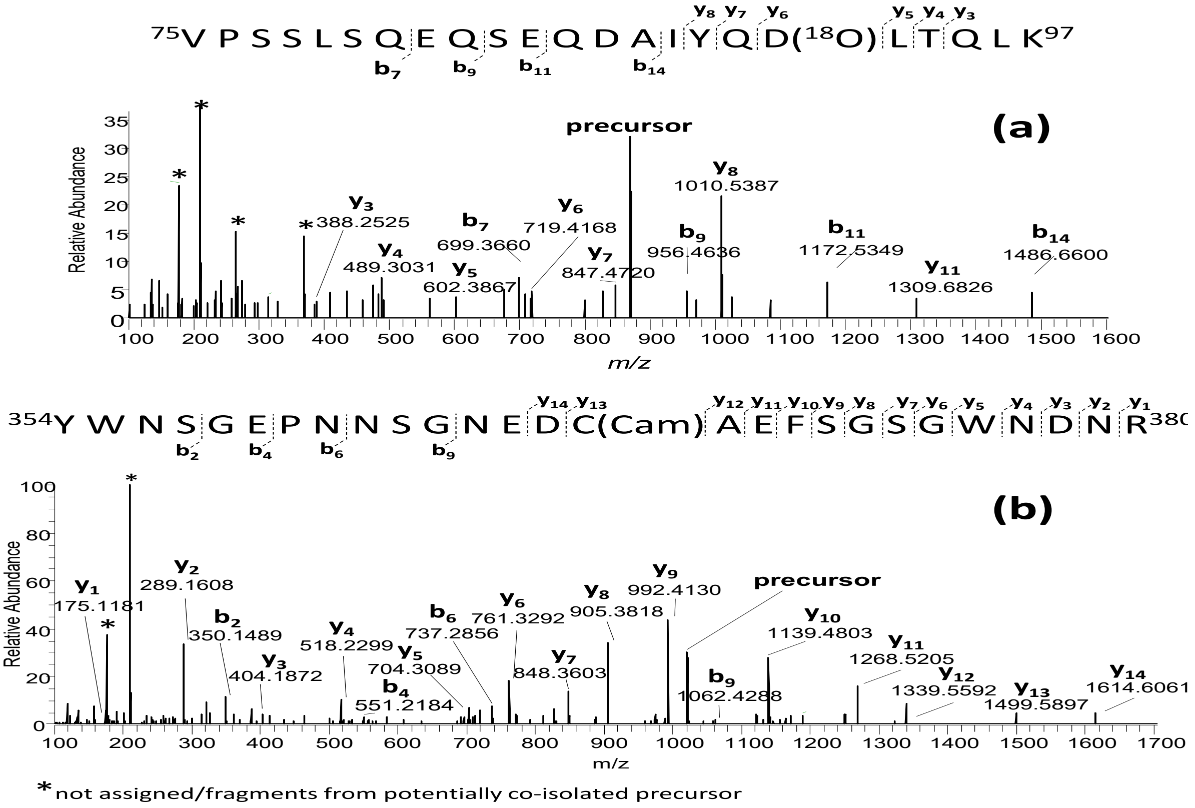

**Materials and Methods:**

**Plasmids, antibodies and shRNAs**: CD209 cDNA (accession# BC110615), CD209L (accession# BC038851) and ACE2 (accession # BC039902.1) were cloned into a retroviral pQCXIP vector. Retroviruses were produced in 293-GPG packaging cells as described (1) and viruses were used to express CD209L, CD209 and ACE2 in HEK-293 cells. Soluble CD209L (amino acids 71-390) and Fc-CoV-2-S-RBD (Accession # MN975262.1, amino acids 319-514) constructs were generated by PCR amplification and cloned in frame with a human Fc fragment of IgE (accession # BC005912) and cloned into a pQCXIP-Myc vector and expressed in HEK-293 cells. SARS-CoV-2-S-RBD-HIS (GenBank: MN975262.1) was cloned into pVRC vector containing a HRV 3C-cleavable C-terminal SBP-His8X tag). SARS-CoV-2-S-RBD was produced by transient transfection in mammalian Expi293F suspension cells. Supernatants were harvested five days post-transfection and passed over Cobalt-TALON resin (Takara) followed by size exclusion chromatography on Superdex 200 Increase 10/300 GL (GE Healthcare) in PBS. Purity was assessed by SDS-PAGE analysis. The Lai-lucΔenv (Env-deficient HIV-1 containing a luciferase reporter gene in place of Nef), has been described previously (1). The lentiviral vector pHAGE-Nanoluciferase was generated via PCR amplification of the nanoluc ORF with the following primers: 5'-ATTGCGGCCGCCATGGTCTTCACACTCGAAGATTTCG-3' and 5'-TGAGGATCCTTACGCCAGAATGCGTTCGCAC-3'. The PCR amplified Nanoluc orf was subsequently cloned into the pHAGE-Zsgreen lentiviral vector (a gracious gift of Dr. Darrell Kotton, BU CReM) using NcoI and BamHI restriction enzymes. The lentiviral packaging plasmid, psPAX2 has been described previously (2). The SARS-CoV-2 S/gp41 expression plasmid (a gracious gift of Dr. Nir Hachoen, Broad Institute) expresses a codon-optimized version of SARS-CoV-2 S and was modified to include the eight most membrane-proximal residues of the HIV-1 envelope glycoprotein cytoplasmic domain after residue 1246 of the S protein. Anti-ACE2 antibody (Cat # 4355) and polyclonal anti-CD209 antibody (Cat # 13193) were purchased from Cell Signaling Technology (Danvers, MA). Polyclonal anti-CD209L antibody (Cat# AV42396, Sigma-Aldrich) was purchased from Sigma-Aldrich. CD209L shRNA (cat# sc-42859-SH) and ACE2 shRNA (cat# sc-41400-SH) were purchased from Santa Cruz Biotechnology. The shRNA Plasmids used in this study are a pool of three to five lentiviral vector plasmids which each encode a target specific 19-25 nt shRNA with a 6 bp loop.

**Infection of HUVEC-TERT cells and immunofluorescence analysis:** HUVEC-TERT cells (1 x $10^5$) were seeded in 8-well chamber slides. The next day, the cells were infected with SARS-CoV-2 at the indicated multiplicity of infection (MOI). One day post infection, the cells were fixed in 10% neutral buffered formalin for at least six hours at 4 °C and were then removed from the BSL-4 laboratory for staining and imaging analysis. In brief, the cells were permeabilized with acetone-methanol solution (1:1) for five min at -20 °C, incubated in 0.1 M glycine for 10 min at room temperature and subsequently incubated in blocking reagent (2% bovine serum albumin, 0.2% Tween 20, 3% glycerin, and 0.05% $NaN_3$ in PBS) for 20 min at room temperature. After each step, the cells were washed three times in PBS. The cells were incubated for one hour at room temperature with a rabbit antibody directed against the SARS-CoV nucleoprotein (Rockland; 1:1000 dilution in blocking reagent), which also cross-reacts with the SARS-CoV-2 nucleoprotein (3). The cells were washed four times in PBS and incubated with goat anti-rabbit antibody conjugated with AlexaFluor488 for one hour at room temperature (Invitrogen; 1:200 dilution in blocking reagent). 4',6-diamidino-2-phenylindole (DAPI; Sigma-Aldrich) was used at 200 ng/ml for nuclei staining. Images were acquired using a Nikon Eclipse Ti2 microscope with Photometrics Prime BSI camera and NIS Elements AR software.

**S Pseudotyped lenti virus production and viral entry assay**: Single round nanoluc-expressing lentivirus vectors were generated via transient co-transfection of HEK293T cells with pHAGE-Nanoluc, psPAX2 and SARS COV2 S/gp41 plasmids using calcium chloride and BES buffered saline (2). Alternatively, HEK293T cells were co-transfected with Lai-lucΔenv and SARS-CoV-2 S/gp41 plasmids for generation of luciferase-expressing single-cycle lentiviruses. Virus containing cell supernatants were harvested two days post transfection and filtered using a 0.45-µm syringe filters, aliquoted and stored at -80 °C until further use (2). The $p24^{gag}$ content of the virus stocks was quantified using a $p24^{gag}$ ELISA, as described before (2).

**S Pseudotyped lentivirus Infections**: HEK293, HEK293-CD209L, HEK293-CD209 and ACE2/HEK-293 cells ($2 \times 10^4$/well) were infected by spinoculation, as described before (2). Cells were lysed 48 hours post infection, and cell lysates were quantified for firefly luciferase or nanoluciferase activity per the manufacturer's instruction (Promega).

**CD209L neutralizing assay**: HUVEC-TERT cells were seeded ($2 \text{ X } 10^4$/well) in a 96-well plate (Corning) for overnight. On the next day, cells were media was removed and replaced with control conditioned medium (CM) from HEK-293 cells or CM from HEK-293 cells expressing sCD209L (approximate concentration 1.5µg/ml) followed by infection of cells with mock or SARS-CoV-2 pseudovirus (5ng/well). After 24 h, cells were washed twice and followed by the addition of 50 µL Nano luciferase substrate (Promega), incubated at room temperature for 5min and luciferase signals were measured using a microplate reader (BioTek) as per the manufacturer's instructions.

**Immunofluorescence staining of human tissues**: Paraffin-embedded sections of normal human organs were obtained from the Department of Pathology archive without any personal identifiers. Tissues were sectioned into 5-µm sections and stained. They were processed for immunofluorescence assay using the following antibodies: Rabbit polyclonal anti-CD209L (1:100) dilution, Rabbit polyclonal anti-CD209 (1:100) dilution), anti-mouse CD31 (Abcam, ab9498, 1:100 dilution), anti-mouse MUC1 (Abcam, ab70745, 1:50 dilution), Aquaporin 1 anti-mouse (Abcam, ab9566, 1:50 dilution). Rabbit polyclonal antibody (Abcam, ab37415), and appropriate isotype mouse IgG1 and IgG2 control antibodies were used.

**Far-Western blotting**: Cell lysate derived from HEK-293 cells expressing CD209L-Myc were incubated with BSA or HIS-STRP tag S-RBD. HIS-STRP tag S-RBD bound proteins were purified with immobilized NeutrAvidin gel (Pierce Inc.) according to the manufacturer's instructions. Proteins were eluted from the NeutrAvidin gel and resolved in SDS-PAGE and the membranes were blotted with an anti-Myc antibody.

**Dot blot assay**: Dot blots were carried out on the dot blot apparatuses according to the manufacturer's instructions. Briefly, a protein of interest was blotted on the PVDF membrane as indicated in the figure legends followed by standard Western blotting technique. For dot blot analyses described in Figure 5B, 25 µg of whole cell lysate from HEK293/CD209L was spotted onto the PVDF membrane after activation with methanol using a vacuum system. The membranes were treated with 20 µL PNGase F (500 units/µL) or with 20 µL Endo H (500 units/µL) (New England Biolabs) in 500 µL water at 37 °C for 4 h. Membranes were washed three times (5 minutes each) with Western Rinse buffer (20 mM Tris and 150 mM NaCl) followed by incubation with S-RBD-HIS (1µg/ml) for 1 h. The proteins were visualized using streptavidin–horseradish peroxidase–conjugated secondary antibody via chemiluminescence system. Image J software, an open source image-processing program, was used to quantify the blots.

**Cell culture and cell lines**: Vero E6 cells (ATCC CRL-1586), HEK293T cells, HEK-293 cells, HEK-293 cells expressing various constructs and HUVEC-TERT cells were maintained in Dulbecco's modified Eagle medium (DMEM) supplemented with 10% fetal bovine serum (FBS), L-glutamine (2 mM), penicillin (50 units/ml) and streptomycin (50 mg/ml). Primary human umbilical vein endothelial cells (purchased from ATCC) were immortalized by pBABE-TERT-hygro (Plasmid #1773, Addgene) and are hereafter named HUVEC-TERT.

**SARS-CoV-2 virus propagation**: SARS-CoV-2 isolate USA_WA1/2020 was kindly provided by Natalie Thornburg (Principal Investigator, CDC) and the World Reference Center for Emerging Viruses and Arboviruses (WRCEVA). SARS-CoV-2 stocks were grown in Vero E6 cells and virus titers were determined by tissue culture infectious dose 50 (TCID50) assays. All work with SARS-COV-2 was performed in the BSL-4 facility of the National Emerging Infectious Diseases Laboratories (NEIDL) at Boston University, following approved SOPs.

**Mass Spectrometry analyses**: For mass spectrometry analyses, myc-tagged CD209L was immunoprecipitated from HEK/CD209L lysates using Pierce Anti-c-Myc magnetic beads (Thermo Fisher Scientific, Waltham, MA, USA) according to the manufacturer's protocol. Following elution, CD209L was treated with sequencing grade modified trypsin (Thermo Fisher Scientific, Waltham, MA, USA) at an enzyme-to-protein ratio of 1:20. Following the digestion with trypsin, half of the sample was additionally digested with endoproteinase AspN (New England Biolabs, Ipswich, MA, USA) an enzyme-to-protein ratio of 1:20, according to the supplier's protocol. The solutions containing the peptides from the tryptic and trypsin-AspN digests were then divided in half. Half of each sample was treated with PNGase F (New England Biolabs, Ipswich, MA, USA) in the presence of $H_2{}^{18}O$ (Cambridge Isotope Laboratories, Tewksbury, MA, USA), while the second half of the sample was reserved for glycopeptide analyses. The peptide mixtures were desalted using 0.6 µL $C_{18}$ ZipTips (Millipore Sigma, Burlington, MA, USA), prior to mass spectrometry analysis. nUPLC-MS/MS analyses were performed on an Orbitrap Fusion Lumos Tribrid mass spectrometer (Thermo Scientific) equipped with an ACQUITY UPLC M-Class system (Waters) and a TriVersa NanoMate (Advion). Online peptide trapping and separation were performed for peptide/glycopeptide analyses. For chromatographic separation, a nanoEase Symmetry C18 UPLC Trap Column (100 Å, 5 µm, 180 µm × 20 mm, Waters) was used for trapping, and a nanoEase MZ HSS C18 T3 UPLC Column (100 Å, 1.8 µm, 75 µm × 100 mm, Waters) was used for separation. The peptide trapping step was performed at 4 µL/min for 4 min with 1% acetonitrile and 0.1% formic acid (Solvent A). Following the trapping step, peptides were separated on the analytical column according to the following conditions: 0–1 min: 2% B, 1–3 min: 2–5% B, 3–43 min: 5–40% B (Solvent B: 99% acetonitrile and 0.1% formic acid). All analyses were performed in the positive mode, with the RF lens set to 30%. MS scans were acquired with the following settings: 120,000 resolution @ *m/z* 400, scan range *m/z* 350–2000, 1 µscan/MS, AGC target 1 x $10^6$, and a maximum injection time of 50 ms. For HCD analyses, initial MS2 scans (35% collision energy) were acquired with the following settings: 15,000 resolution @ *m/z* 400, scan range *m/z* 100–2000, 1 µscan/MS, AGC target 1 x $10^6$, and a maximum injection time of 100 ms. Based on the initial HCD (35%) MS2 scans, a second fragmentation step was triggered if at least two glycopeptide oxonium ions were detected (at *m/z* 204.0867, HexNAc; *m/z* 138.0545, HexNAc – $CH_6O_3$; and *m/z*

366.1396, HexNAcHex) within a 15-ppm mass tolerance. This second product ion triggered MS2 (15/25/35% collision energy) was acquired with the following settings: 30,000 resolution @ *m/z* 400, scan range *m/z* 100–2000, 1 µscan/MS, AGC target 1 x $10^6$, and a maximum injection time of 150 ms.
